# The Effects of Depression on The Neural Correlates of Reward Sensitivity in Poverty

**DOI:** 10.1101/2022.07.28.501886

**Authors:** Hiran Perera-W.A., Rozainee Khairuddin, Khazriyati Salehuddin

## Abstract

Existing studies have identified that depression and depressive symptoms are associated with reduced sensitivity to feedback processing, which is a core ability that determines the success of human actions. However, a key individual difference which is the socioeconomic status (SES) has been largely ignored in this field because the recent trend of research has suggested how it relates to various cognitive domain-specific neural systems. Because depression is a widespread mental health condition that is more prevalent among the poor, it can potentially play a role in the association between poverty and feedback processing. With a sample of 80 adults recruited from low to high-income communities, the current study examined the role of depression on the relationship between poverty and feedback processing by using feedback negativity (FN) event-related potential (ERP), which is a well-known ERP component that is indexed by response feedback indicating losses versus gains. Consistent with previous studies, high depressive symptoms were associated with reduced FN amplitude across our sample. SES was negatively associated with FN and depressive symptoms, which indicates reduced reward sensitivity to feedback among the low-SES individuals who are also mostly depressed. However, no association between SES and reward sensitivity was observed when it was controlled for depression. Findings in this study suggest the importance of partial out the variance accounted for by depression when studying responses to reward sensitivity in poverty.

## INTRODUCTION

Our goal-directed behaviors are facilitated by the capacity to monitor and adapt to continuously changing environments. Our brains possess the ability to quickly and efficiently evaluate decisions that we make based on the feedback we receive. Monitoring those feedback-related responses plays a significant role in guiding goal-oriented behaviors. One important element of the feedback-related action monitoring is the sensitivity to rewards, which is the degree to an individual’s behavior is motivated by the reward-relevant stimuli (Carver & White, 1994; Schultz, 2006). Findings suggest that reduced activation of our reward system can lead to reduced sensitivity to rewards (Balconi et al., 2015; Braver et al., 2014), as well as reduced motivational attainment of goal-directed behaviors (Balleine et al., 2007; Braver et al., 2014).

A key individual difference which is the socioeconomic status (SES) has been a contributing factor to the variabilities in reward sensitivity. SES is a construct that most often refers not only to differences in income and wealth but also to subjective positions in society and levels of education (Perera-W.A. et al., 2021). Resource scarcity in poverty exacerbates the tendency to react more strongly to the prospect of losses (Adamkovič & Martončik, 2017). When presented with negative life events such as lack of financial ability, vulnerable individuals often tend to take riskier decisions such as gambling, lack of health care plans, and increased debts, eventually leading to worsening the conditions that they live in (Carvalho et al., 2016; Ong et al., 2019; van der Maas, 2016). Children for instance, who grew up in poverty households on average, experience increased behavioral problems (Dearing et al., n.d.) with increased sensitivity to reward-seeking behaviors (Galván et al., 2013; Lansford et al., 2017). Similarly, with adults, findings suggest an inverse relationship between SES and risk-taking behaviors such as engaging in smoking, alcohol abuse, and unhealthy eating habits (C.-C. Chen & Yin, 2008; E. Chen et al., 2015; Payne et al., 2017).

But how exactly does poverty impacts the sensitivity to rewards? There are two competing hypotheses. Firstly, those who are from poverty households often experience high neighborhood violence, poor nutrition, and low cognitive stimulation during the early developmental period (Evans, 2004; Kim et al., 2016; Noble et al., 2006, 2015) which can largely contribute to the changes in the structural development of cortical and subcortical systems (Noble et al., 2015) that subserve reward sensitivity and reward processing. These systems include the amygdala, basal ganglia, thalamus and hippocampus (Gianaros & Hackman, 2013; Gonzalez et al., 2016; McLaughlin et al., 2014; Nusslock & Miller, 2016; Sheridan & McLaughlin, 2014), which play a crucial role in mediating threats and alter reward processing (Gonzalez et al., 2016). Differences in the sensitivity to rewards can also be attributed to processing inconsistencies in the neurophysiological mechanisms (Potts et al., 2006). Converging evidence across multiple brain systems suggests a dynamic interaction between the prefrontal cortex and subcortical regions, with an interplay of dopamine activation systems (Crone & van der Molen, 2004; Frank & Claus, 2006; Potts et al., 2006) which predicts potential gains/losses and determine how to respond to environmental challenges.

Additionally, cortical structures such as the anterior cingulate circuit (ACC), ventro-striatal neural network, and lateral prefrontal cortex also involves in the anticipation and receipt of reward (Braver et al., 2014; Haber & Knutson, 2010). These structures are collectively involved in the automatic updating of goal representations, cognitive flexibility, and effortful engaging or inhibition processes (Aarts et al., 2013; Cools et al., 2011) and are complexly mediated by the dopaminergic neurotransmission as the biological basis (Cools et al., 2011). When faced with negative feedback, neurophysiologically, a midfrontal negative-going amplitude shift (compared to positive action responses) is observable, which reflects an early binary mechanism of action evaluation (Hajcak et al., 2006).

Secondly, a growing number of evidence suggests a strong negative association between poverty and depression (Heflin & Iceland, 2009; LeMoult et al., 2020; LeMoult & Gotlib, 2019; Lorant, 2003; Patel, 2001). Neural responses to negative feedback and negative processing biases (exaggerated responses to negative performance feedback) have been observed in individuals with depression (Bress et al., 2015; Kring & Bachorowski, 1999). Because depression is associated with increased sensitivity to threats and reduced sensitivity to rewards (Shankman et al., 2007, 2013) there is a profound impact on cognitive processes such as cognitive control, and outcome monitoring (Olvet et al., 2010; Ruchensky et al., 2020; Tang et al., 2013). A considerable amount of work has been devoted to distinguishing reward sensitivity on negative affective states such as stress and depression (Anisman & Matheson, 2005; Berghorst et al., 2013; Keren et al., 2018). A blunted neural response to reward (Keren et al., 2018; W.-N. Zhang et al., 2013) suggests emotional/motivational pathway dysfunctions in depression during reward-related processing. Depressed individuals perhaps assign a negative value to unfavorable outcomes compared to favorable outcomes, and this response is indexed in feedback-related negativity (FRN) event-related component by ACC-related performance monitoring (Mueller et al., 2015; Santesso et al., 2008).

It has been posited that the FRN has a specific relationship to depressive symptoms, specifically over the error-related negativity potential, which is larger among those who are anxious (Bress et al., 2015). An association between depression and blunted FRN has also been reported in adolescents due to their hypersensitivity to negative outcomes and attenuated responsiveness to rewards (Bress et al., 2012; Bress & Hajcak, 2013). However, the opposite trend has also been observed. In a sample of 26 depressed female adolescents, there was an increased FRN response to losses versus for rewards in comparison to the healthy female adolescents (Webb et al., 2017). Findings indicate that the FRN is a potential neural marker for depression (Bress et al., 2015) with a blunted FRN effect in depression. Keren et al. (2018) found a significant combined effect size of *d* =.5 in studies with participants under age 18, which suggests that those who are depressed experienced less differentiation between gain and loss feedback from the frontal lobe responses, exerted potentially as a result of a weaker response to gain feedback (Keren et al., 2018). This effect was not observed among the adults, and it remains unclear how socioeconomic status further influences the reward processing among those who are depressed.

### Feedback-Related Negativity (FRN)

Feedback-related negativity (FRN) ERP component occurs approximately 250ms after feedback and can be observed at fronto-central scalp region (Gehring & Willoughby, 2002). The FRN component is thought to elicit from ACC region, which is the brains action monitoring system, and it is sensitive to reward prediction errors, negative valence, and reward magnitude feedback outcomes (Cohen & Blum, 2002; Gehring & Willoughby, 2002; Holroyd et al., 2006). Typically, negative processing stimuli produce feedback responses to indicate that a negative outcome has happened, such as loss of money or incorrect responses. However, importantly, positive feedback does not elicit a FRN which indicates that the negative and positive feedback-related ERPs are initiated by separate systems (Holroyd et al., 2006; Miltner et al., 1997). FRN effect (the difference between negative and positive outcomes), also known as feedback negativity (FN) is more reflective on the instantaneous outcome (Osinsky et al., 2017) when participants had to choose between two alternatives, which can lead to either a favorable (wins) or unfavorable (losses) outcomes (for example, wins are positive and losses are negative outcomes) (Cui et al., 2013; KreuSSel et al., 2012; Mussel et al., 2015). The FN effect signals the reward prediction error (i.e. when an outcome is worse or better than expected) which can optimize task behaviors and reinforcement learning (Osinsky et al., 2017; Sambrook & Goslin, 2014). It is prominent when there is a strong dissociation between the gain and the loss outcome yet smaller when the perceived difference in the outcome is much lesser. The FRN effect is frequently observed in a simple gambling or a guessing task (Carlson et al., 2011; Foti et al., 2011; Hajcak et al., 2006; Holroyd et al., 2006; Moser & Simons, 2009) when the outcomes are generally risky, random or uncertain. The FN is computed by subtracting the ERP to gains from losses, which results in amplitude with a negative deflection and reward positivity (RewP) by subtracting loss from gain (positive deflection). The FN effect is prominent when there is a strong dissociation between the gain and the loss outcome yet smaller when the perceived difference in the outcome is much lesser. Because FRN has been correlated with depression and no studies have examined its association with SES (Perera-W.A. et al., 2021), we examined FN to evaluate the extent to which poverty and depression are drawn towards reward processing.

### Current study

The goal of the present study was to examine the association between socioeconomic status and FN responses to positive and negative unexpected feedback outcomes and to see how subclinical depression influence this relationship. The use of depression was justified by the fact that low-SES and depression typically co-occur, which was also found to exert its effects on the neural responses to error monitoring. Furthermore, SES is strongly related to variabilities in reward sensitivity and high depression levels, which are often seen among low-SES.

This study employed an experimental paradigm of the reward guessing task which was a simplified version of the widely used Iowa Gambling Task (IGT) (Bechara et al., 1994) to elicit an unambiguous measure of reward sensitivity. IGT is an excellent tool for examining decision uncertainties. The task simulates uncertain real-life decision-making of gains and losses where subjects had to pick their responses from a choice of 4 decks of cards; two out of four decks consist of large wins or large losses, while the remaining typically consist of smaller wins or loses (Bechara et al., 1994; Toplak et al., 2010). Participants chose these options without having an explicit knowledge about which deck will lead to greater reward or greater losses, which means that the decision-making was exclusively based on the knowledge of the risk assessment associated with each deck (Bechara et al., 1994; Brand et al., 2007). Several variations of the IGT have been used to see response choices between healthy participants and VMPFC-damaged patients (Bechara et al., 1994), and more recently, gender differences in reward sensitivity (Garrido-Chaves et al., 2021).

In each trial, participants chose between one of two rewards options which subsequently could result in a Win or Lose outcome. We administered this task to a non-clinical sample of adults from low to high SES backgrounds in order to investigate the relationship between SES and neurophysiological response to rewards (indexed by the FN ERP component). We predicted that consistent with findings from behavioral and neuroimaging studies, the FN would differentiate monetary losses from monetary gains and that the difference between loss and gain trials would be attenuated among the low-SES with higher depressive symptoms scores. We expected that this relationship might specifically be driven by the neural response to losses. Furthermore, based on the findings, we predicted that reward sensitivity, as reflected by the FN difference, would be different when it is controlled for depression.

## METHODS

### Participants Selection

A total of 80 (out of 88) right-handed participants living in the urban area of Kuala Lumpur, Malaysia, participated in this study. Low-income participants were recruited through flyer distribution in several low-income housing communities. We also used social media and snowballing sampling method to recruit participants from various income backgrounds. All participants are married and live in households that share household commodities with other family members. Prior to their participation, participants were screened for chronic mental health issues, traumatic brain injuries, and any alcohol dependencies. All participants gave their written informed consent before starting the study.

We excluded data from eight participants due to high EEG artifacts (70 percent of EEG epoched data had peak-to-peak amplitude greater than 100*µ*V). All the remaining participants (*n* = 80, 67.6% are females) were from the three major ethnic groups that make up the Malaysian population, namely Malay, Chinese, and Indian ethnicities. The mean age of the remaining participants was 40.2 years old. (range = 42, *SD* = 11.8). Participants received a maximum of RM 60 (USD 14 equivalent) upon successfully completing the study.

The study was conducted at the community room of the Ministry of Women, Family, and Community Development (Kementerian Pembangunan Wanita, Keluarga dan Masyarakat, KPWKM), in Kuala Lumpur, Malaysia. Individuals who were interested in participating were given a time slot and a pre-confirmed date. Upon arrival, participants were given the informed consent form, a financial background questionnaire, and a depression scale (CES-D) which was then followed by the preparation for the EEG recording. This research received approval from the University Kebangsaan Malaysia (UKM) ethics committee (UKM PPI/111/8/JEP-2018-339) and was funded by the IDEA UKM through the Dana Cabaran Perdana grant (DCP-2017-014/1).

### Instruments

#### Depression Measures

To measure common depressive symptoms of the participants, this study utilized the Malay-translated version by (Mazlan & Ahmad, 2014) of the The *Center for Epidemiological Studies-Depression* (CES-D) scale (Radloff, 1977). The Cronbach’s alpha of the translated instrument was 0.75, with test-retest reliability of the total score of *r* = .69. The instrument consists of a 20-item scale, which measured depressive symptoms, including mood, helplessness, loss of energy, sleep and appetite issues participants experienced within the last week. Each item was worded as a self-statement, where participants had to indicate how often they felt that way in the past week. The scoring ranged from 0 to 60 and 16 or greater was the cut-off point for depression. This self-reported scale has been widely used in depression research and have been deemed appropriate to measure depressive symptoms among non-clinical samples (Amtmann et al., 2014; Shafer, 2006).

#### Monthly Income Measures

Due to the complex financial resources and capital attached with SES, it is important to properly conceptualize and operationalize to assess the socioeconomic context (see Perera.W.A. et al 2021). Apart from household income, which is the most common variable associated with when operationalizing SES, there are several other indicators such as parental education and occupation have also been considered, yet these variables can have different implications. For instance, household income and parental education can predict academic success (Duncan and Magnuson, 2003).

In order to facilitate the identification of participants at or near national poverty cutoff line (in Malaysia), we made an effort to recruit a sample across the broader SES spectrum. All participants reported their monthly household income on a 9-point Likert scale which provided a fine-grained breakdown of income at the low and high levels (e.g., 1 = RM1000 or less, 2 = RM1001 – RM2000, 3 = RM2001 – RM3000, etc.). Two broad income categories (low and high-SES) were used in the behavioral outcome analysis. The continuous 9-point scale SES variable represents the full range of income used for all remaining analyses. This representation of participants across the full spectrum of income allowed for a robust test of the effects of income on cognitive control.

### Experimental Paradigm

In this study, a variation of the Iowa Gambling Task (IGT) was developed and administered to the participants using Psychopy version 3.5 software (Peirce et al., 2019) on an Acer 21” All-In-One desktop PC. Participants responded with an attached external QWERTY keyboard. The experimenter was present throughout the testing session; however, no feedback was provided to the participants when they performed the task. Each participant sat approximately 75-cm away from the computer screen with the keyboard placed in front.

The task consisted of 160 trials, which were presented in four blocks of 40 trials separated by intervals that lasted a maximum of 3-minutes. Because participants in this study consisted of poverty and low-SES communities, and because the FRN-ERP component was found to be less sensitive to magnitude rather than valence (Cui et al., 2013), we simplified the task by removing the magnitude option. The schematic flow of the task can be seen in Figure 1.

**Figure 1:**
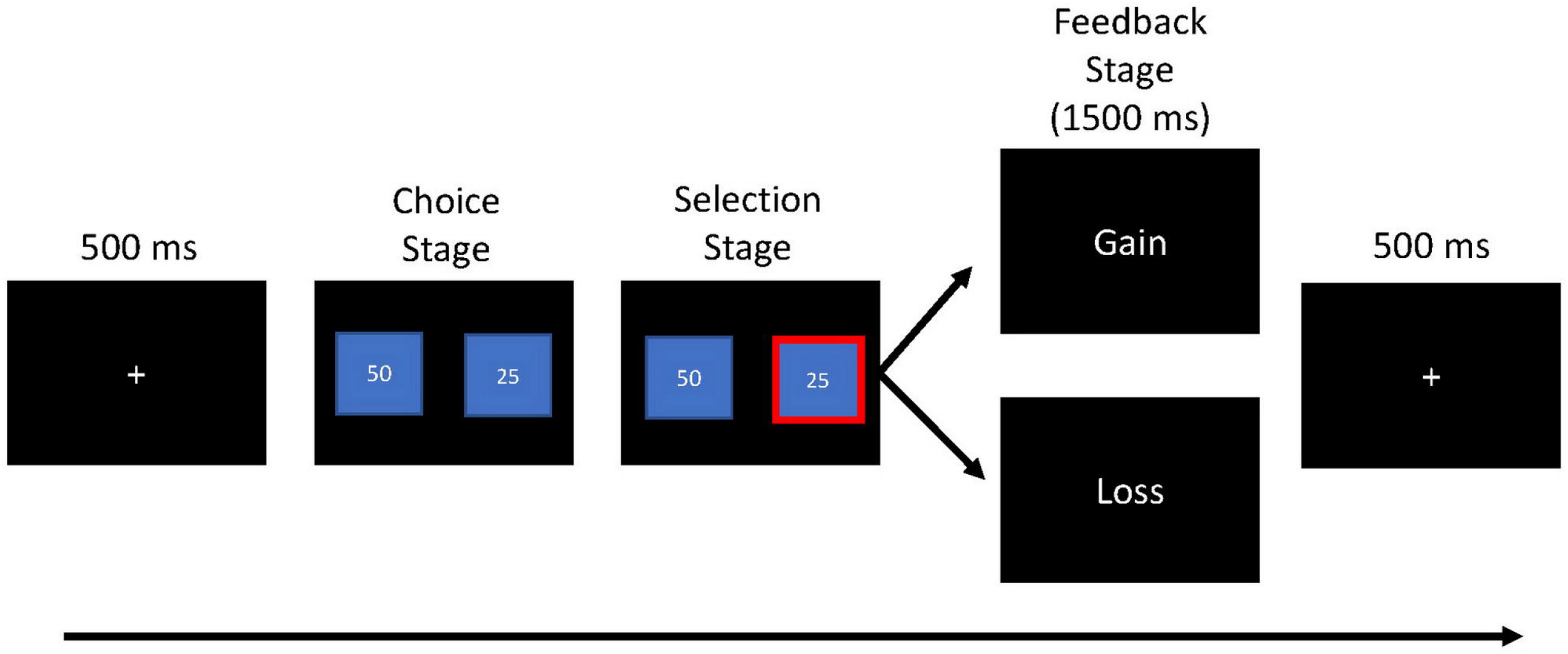
The schematic representation of the modified IGT, which consists of choice stage, selection stage and the feedback stage.

At the beginning of each trial, participants were presented with two monetary options, large (RM50) and small (RM25), left and right on the computer screen. They were instructed to choose one of the options via a keypress, “z” key for the left option, or “m” key for the right option. Participants were instructed that they could either win or lose the actual money on each trial, and the goal was to correctly select the option that would maximize their chances of winning at the end of the experiment.

The task began with a 500ms fixation cross, which was then followed by the choice stage with two monetary choice options. The two options remained on the screen until the participants made a selection. In the selection stage, the fixation cross disappears, and after the response onset, the selected choice was highlighted with a red frame for 1000ms. The selection stage was then followed by the feedback stage indicating the wins or loses, which was presented for 1500ms. Participants were informed that they could either gain (50) or lose (25) during their selection. These values were chosen based on previous similar studies (Tversky & Kahneman, 1982, 1992) to equalize subjective values of gains and losses. A gain was indicated on the feedback screen by the words “You Won”, and a loss was indicated by the words “You Lose”. The probability of winning and losing was precisely 50%; 20 gains and 20 losses feedback trials were presented in fully random order.

### Psychophysiological Recording, Data Reduction, and Analysis

ERP recordings were acquired using Cognionics HD-72 high impedance dry and mobile 32-channel wireless EEG headset at a rate of 500 Hz. Impedance level was maintained at less than 100 Kilo Ohms (Chi et al., 2013). Electrodes were arranged according to the 10/20 system, including FCz. Left and right mastoid electrodes served as ground and reference, respectively (Figure 2) (Perera-W.A., 2022).

**Figure 2:**
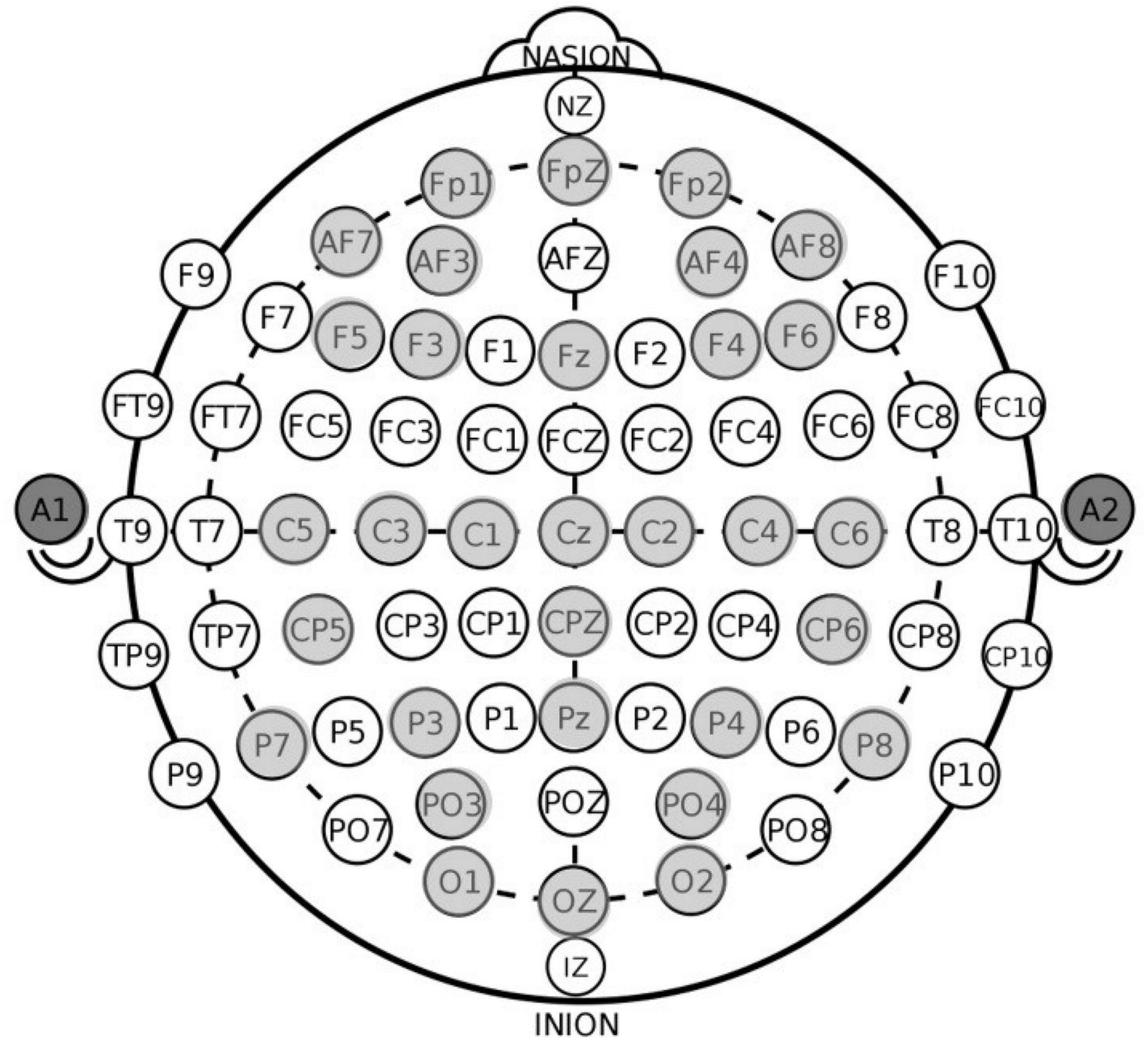
Cognionics 32-channel EEG electrodes positions (marked in grey color). A1 and A2 illustrate ground and reference electrodes, respectively.

The Cognionics system is equipped with an active ground system and a Faraday-cage-like enclosure to prevent interference from electrical noise (Chi et al., 2013; Mullen et al., 2015). This system is ideal for EEG data collection taking place outside of a lab environment. Raw EEG data was converted offline to an average reference. EEG data were pre-processed using 14.1.1b (Delorme & Makeig, 2004) and ERPLAB version 6.1.4 (Lopez-Calderon & Luck, 2014). Data was filtered offline (1 - 30 Hz) and segmented into epochs of 800 ms for the feedback stage, which consisted of 200 ms before and 600 ms after the feedback onset. All epochs were independently and visually inspected for possible artifacts, and the baseline corrected with respect to the pre-stimulus period.

Independent Component Analysis (ICA) was performed on epoched data using the “Infomax” ICA decomposition method implemented in the “*runica()*” function of EEGLAB (Delorme & Makeig, 2004). We next followed standard guidelines (Jung et al., 2000) to detect and remove components accounting for ocular and muscular artifacts. Next, we rejected data epochs with max-min amplitudes exceeding 100µV in 1000ms segments isolated in steps of 100ms. In line with recommended practice, we excluded subjects from the sample who had more than one-third of their trials rejected according to these criteria.

Stimulus-locked ERPs were averaged independently for gains and losses. The ERP activity was quantified as the activity at FCz electrode site, as the average activity from 250ms to 350ms after feedback onset. The feedback negativity (FN) is the activity on losses minus the activity on gains, which has been used in a number of studies (Bress et al., 2012; Crane et al., 2021; Hajcak et al., 2006; Yeung et al., 2005). Here, the difference score provided a measure of neural sensitivity to outcome valance regardless of the source (Bress & Hajcak, 2013; Christie & Tata, 2009; Dunning & Hajcak, 2007; Foti & Hajcak, 2010). Although this scoring method did not allow for investigating the individual contributions of responses to gains and losses and perhaps it can indicate an ERP component where one does not exist (Luck, 2014). On average, 3.01 channels were found to be artifactual. They were interpolated either through spherical spline interpolation or nearest-neighbor replacement.

## RESULTS

### Poverty and Depression Scores

First, we performed a bivariate correlation between SES (utilized as a continuous variable (from low to high-SES) and the level of depression (CES-D). There was a strong negative relationship between variables SES and the level of depression (*r* = -.45, *p* < .001), which indicated that depressive symptoms were strongly associated with low-SES (Figure 3). The income was treated as a continuous variable in most of the statistical analyses to minimize information loss (MacCallum et al., 2002). CES-D score did not correlate with age (*p* > .05).

**Figure 3:**
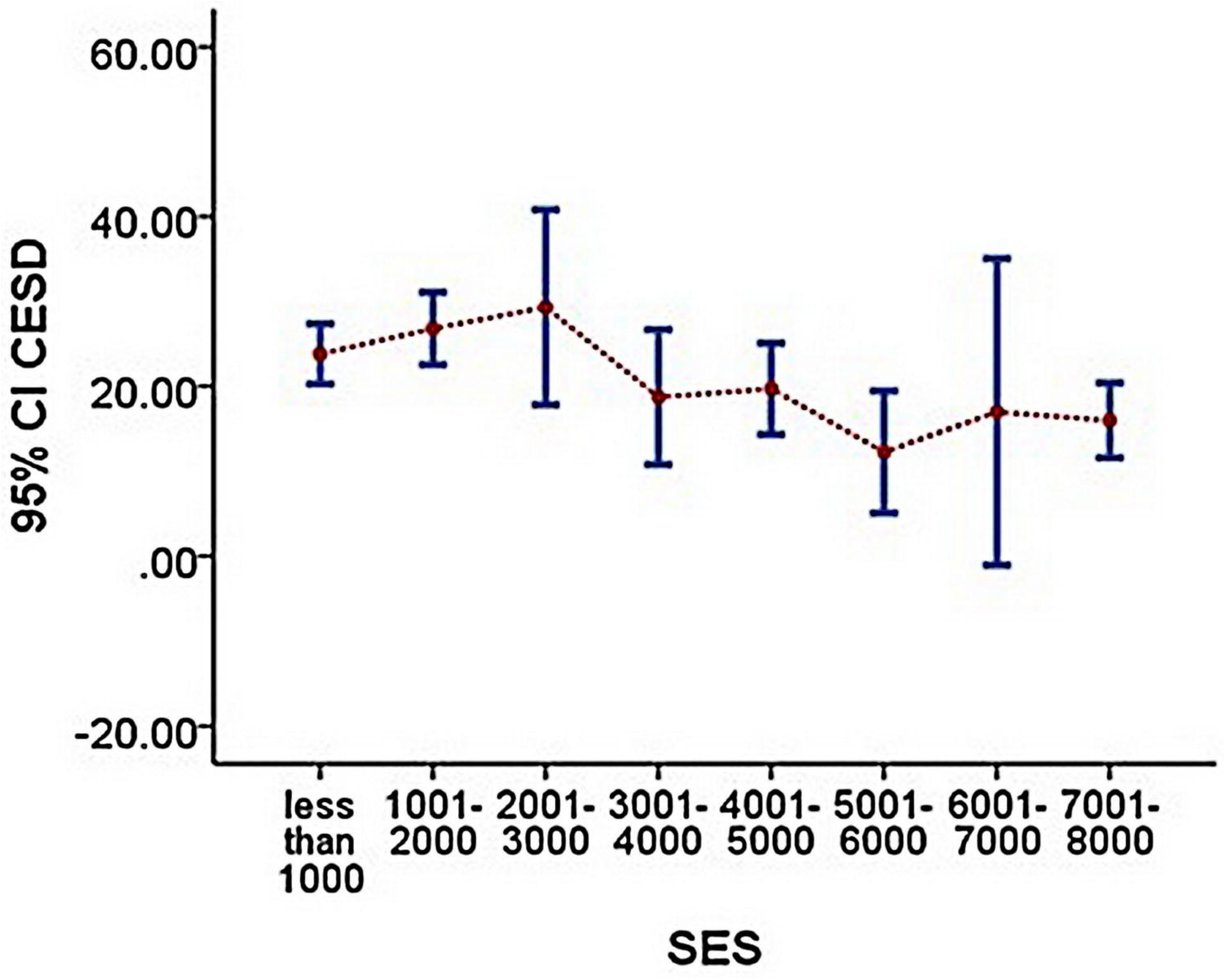
Error bars (95%) indicate the relationship between SES and depression (CESD) scores (p < .001).

Second, to identify whether there was a difference in CES-D scores between SES groups separately (low-SES and high-SES as the grouping variables), an independent samples *t*-test was performed. We found a significant difference in depression scores between low and high-SES groups, *t*(78) = 4.84, *p* < 0.05, 95% CI [5.57, 13.36], where low-SES had a high level of depression compared to the high-SES participants. Males and females had similar depression scores (*p* > .05) in both SES groups. Means, standard deviations, and ranges for CES-D scores are reported in Table 1. Figure 4 represents the box plot of CES-D mean scores between low and high-SES groups for a visual comparison.

**Figure 4:**
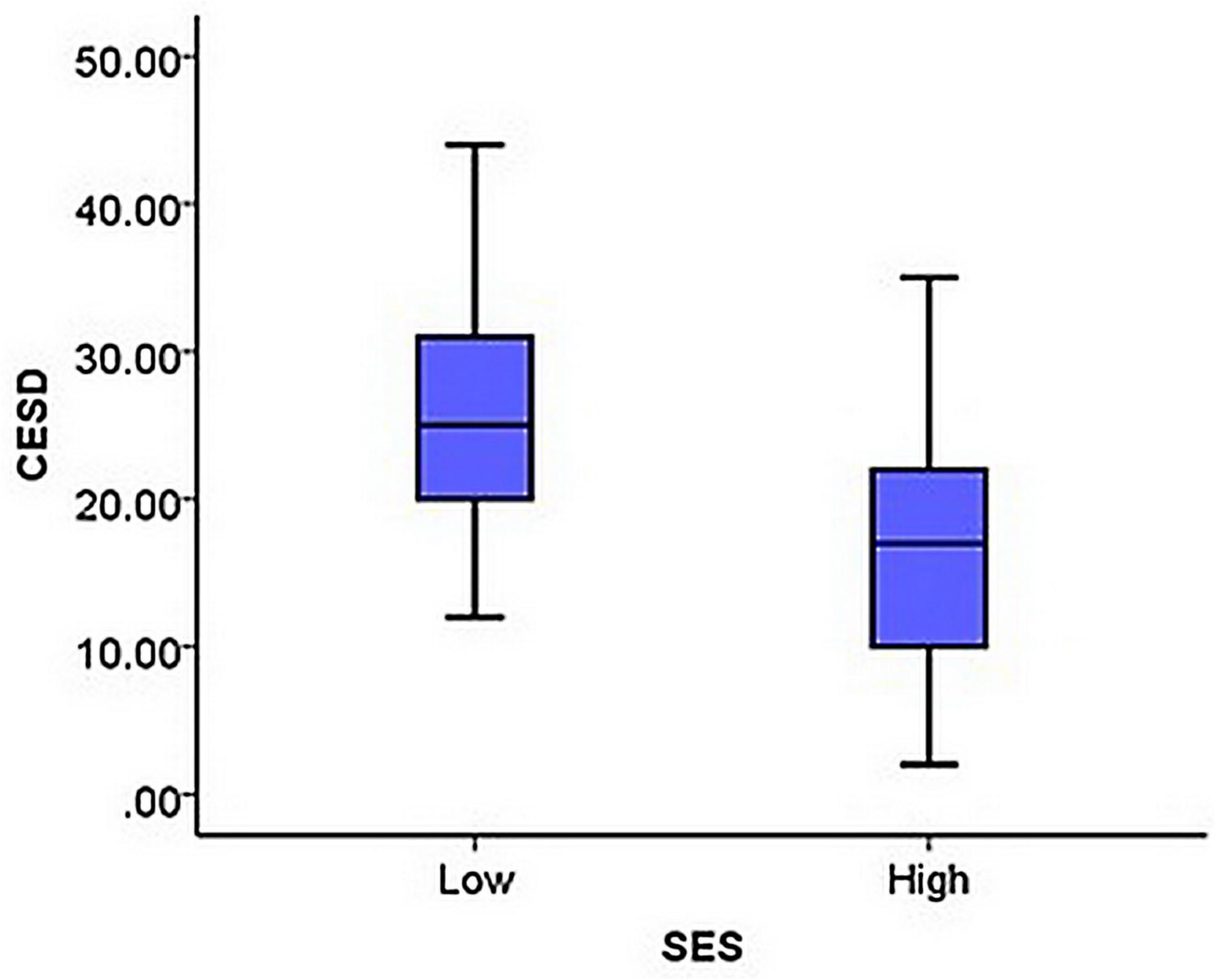
Boxplot Error bars (95%) represent the depression (CESD) mean scores of low and high SES groups

**Table 1:**
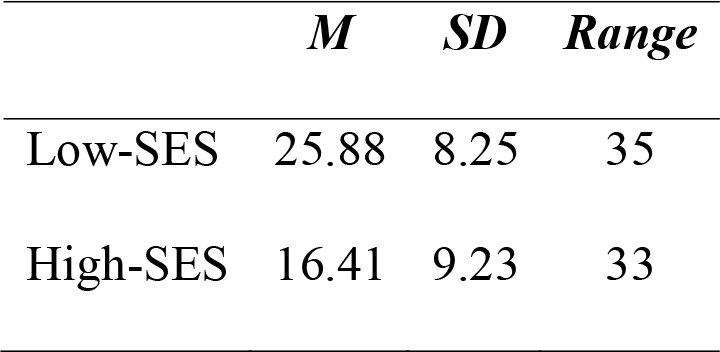
SES and Depression scores (CESD)

|  | *M* | *SD* | *Range* |
| --- | --- | --- | --- |
| Low-SES | 25.88 | 8.25 | 35 |
| High-SES | 16.41 | 9.23 | 33 |

### SES and Feedback Negativity (FN)

Consistent with previous studies (Foti et al. 2011b; Bress & Hajcak 2013), the feedback-locked ERPs were more negative at FCz electrode site. The feedback indicating monetary loss was associated with more negativity at around 350ms compared to feedback indicating monetary gain (Figure 5a). The difference between losses and gains (dFN) was maximal at the frontocentral site.

**Figure 5:**
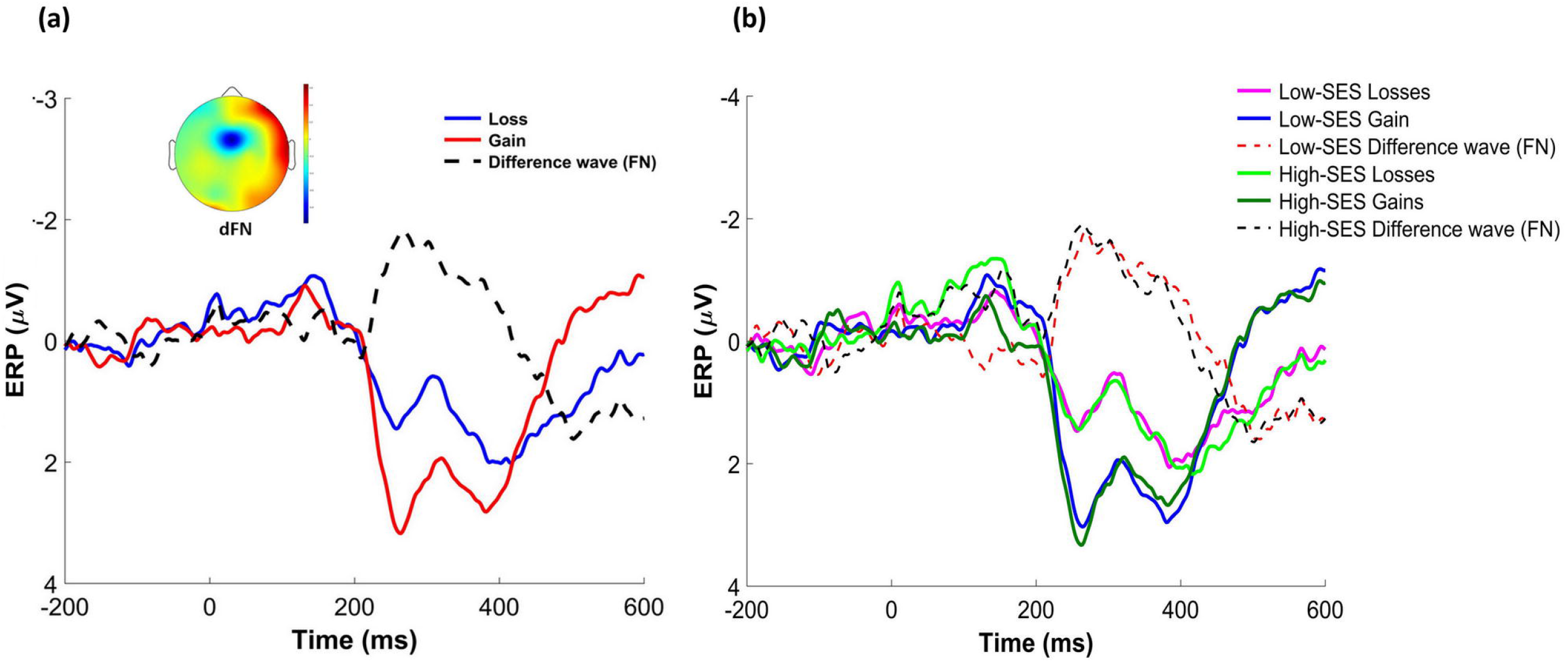
Stimulus-locked event-related potentials to feedback indicating monetary loss (blue) and gain (red), as well as the difference waveform for loss minus gains trials (dashed). Scalp distribution of the difference between loss and gain trials from 250 to 350 ms following feedback onset (a). Low and High-SES-related stimulus-locked event-related potentials to feedback indicating monetary loss and gain, as well as the difference waveform for loss minus gains trials (b).

A paired-samples *t*-test confirmed that monetary loss (*M* = 1.03 µV, *SD* = 2.72 µV) was associated with greater negativity compared to monetary gains (*M* = 2.46 µV, SD = 2.87 µV) which is significant (*t*(79) = −4.41, *p* < .001; Cohen’s *d* = .51). Figure 5b represents FN amplitudes of all gain, losses as well as the difference waves for both low and high SES groups independently.

Series of bivariate correlations were performed, taking SES as a continuous variable. Strong associations were found between dFN amplitude and SES (*r* = −.23, *p* < .05; see Figure 6a), and dFN amplitudes and CES-D (*r* = .22, *p* < .05; see Figure 6b). Table 2 indicates zero-order correlations between dFN and study variables.

**Figure 6:**
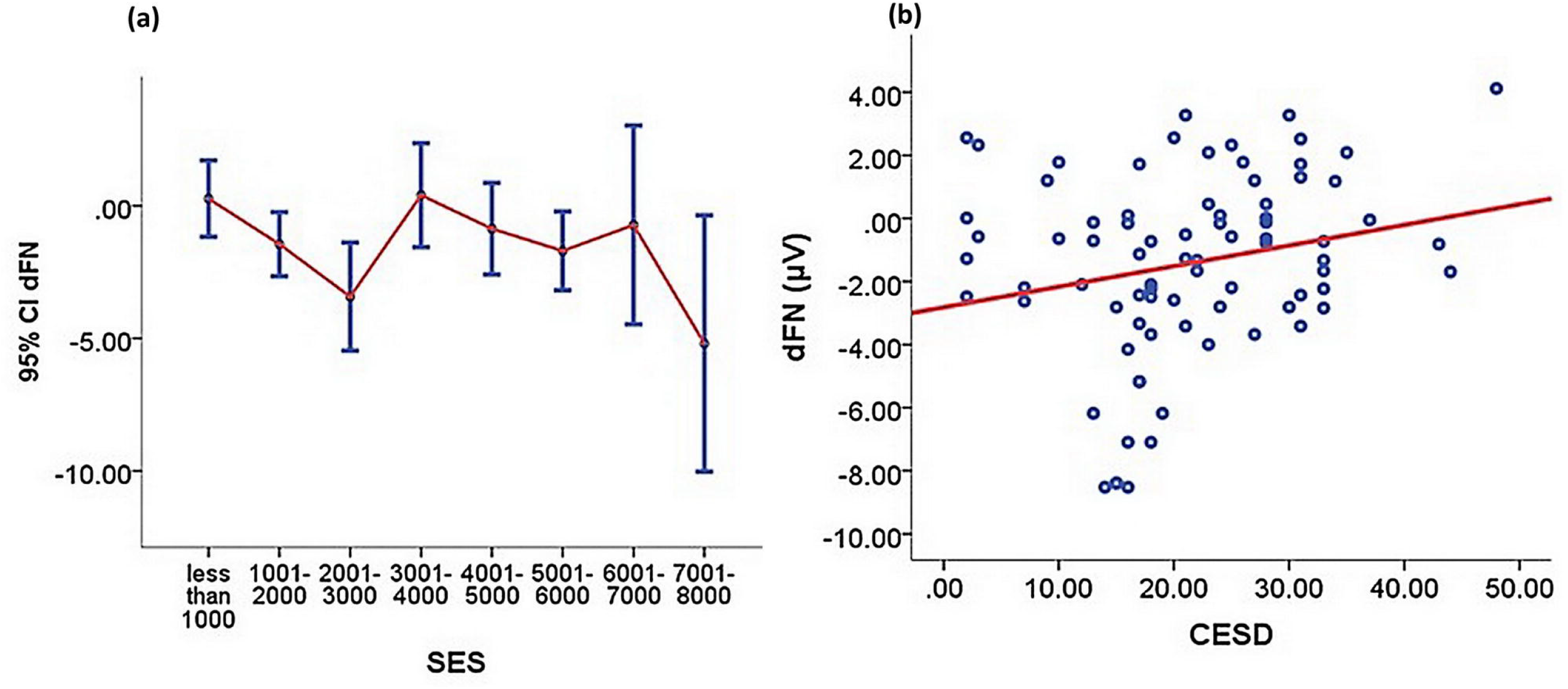
Negative correlation between FN (µV) and SES (p < .05) (a). Scatter plot of feedback negativity (FN: calculated as a difference score between loss and gain trials) and CESD depression scores (p < .05) (b).

**Table 2:** Zero-order correlations between variables

|  | **1** | **2** | **3** | **4** |
| --- | --- | --- | --- | --- |
| SES | - |  |  |  |
| CES-D | -0.45** | 1 |  |  |
| dFN | -0.23* | .22* | 1 |  |
| Loss | -0.10 | 0.01 | .48** | 1 |
| Gain | 0.13 | -0.21* | -.56** | .47** |

\* $p < .05$ ; \*\* $p < .01$ ; \*\*\* $p < .001$

Since dFN difference score is numerically negative, negative correlations indicate that the lower SES spectrum is associated with reduced (more towards positive) dFN amplitude (i.e., reduced neural differentiation between losses and gains, see Figure 6a). Additionally, a positive correlation between dFN and CES-D scores indicates a reduced dFN amplitude (see Figure 6b) among depressed.

Additionally, a partial correlation was performed to determine the relationship between SES and participant’s neural responses to dFN while controlling for depressive symptoms (i.e., partial out the variance accounted for by depression). However, we did not find a significant correlation between SES and dFN amplitude while controlling for CES-D (*r* = −.15, *p* > .05). Findings indicate that depressive symptoms have an influence in the relationship between SES and participant’s neural responses to reward sensitivity, dFN.

## DISCUSSION

The current study examined the neural responses of SES to sensitivity to reward feedback by using feedback negativity (FN) ERP. FN was calculated as the difference between losses and gains, which was maximum at the frontocentral scalp site. We found that overall sensitivity to reward was not significantly different between low and high-SES groups. However, FN amplitude was more negative for losses than gains, and this finding is in line with past studies of gains versus losses characterized by enhanced negativity of FN amplitude in IGT paradigms (Dunning & Hajcak, 2007; Foti et al., 2011; Foti & Hajcak, 2010).

We also examined the influence of depression on the sensitivity to reward. We observed a blunted FN among the more depressed participants. This effect was also consistent with the existing studies (Foti & Hajcak, 2010). However, when losses and gains were considered independently, increased depressive symptoms were negatively associated with gains, but not with losses. When depression was controlled, however, we found that participants’ reward processing was driven by their pathological state and not entirely by their socioeconomic status. This outcome is not entirely surprising given that a large body of existing research suggests that poverty is associated with maternal depression (Eamon & Zuehl, 2001; Garg et al., 2015; Heflin & Iceland, 2009; Tomarken et al., 2004), depressive symptoms in adults (Lorant, 2003), and in children (Flouri et al., 2017; LeMoult et al., 2020; LeMoult & Gotlib, 2019). Several possible justifications and implications for the findings are highlighted in the following sections.

While some researchers have identified that poverty is associated with increased sensitivity to threats (E. Chen & Matthews, 2001; Javanbakht et al., 2015; Kim et al., 2013; Muscatell et al., 2012), there are also studies that suggest that poverty is associated with greater anticipation to rewards (Gonzalez et al., 2016; Romens et al., 2015). There are also limited studies investigating reduced sensitivity to rewards (see review by Yaple & Yu (2020)) in children due to overexposure to conflicting and adverse environments during their developmental period. One possible explanation for the reduced sensitivity to rewards is due to the external stressors associated with poverty and being vigilant to punishment cues when growing up in poverty, which consequently reduce the ability to deal effectively with stressors (Dufford et al., 2020; Evans, 2004; Gonzalez et al., 2016; Kuruvilla & Jacob, 2007; Letourneau et al., 2013). This increases the need for engaging in more riskier and impulsive behaviors due to the heightened sensitivity to rewards (Ellis et al., 2009), by favoring anticipation of rewards system, and suppressing the reward regulatory system (Romens et al., 2015). In our attempt to identify the effects of SES on reward processing, we found that subclinical depression associated with SES plays an important role in guiding the sensitivity to rewards.

The link between subclinical depression and socioeconomic inequality is well-established, which is a risk factor for chronic mental health problems. Stress and depression are most prevalent among the poor (McLoyd, 1998; Ridley et al., 2020). It is evident that not only low-SES reportedly experience depression 1.81 times higher than those who are from the high-SES spectrum (Lorant, 2003), but increasing evidence also suggests that depression is negatively correlated with income (Lund et al., 2011; Ridley et al., 2020). Consequently, the effects of depression, such as hopelessness, sadness, and anxiety, have a significant impact on reward-related processing (Lam et al., 2014). Furthermore, anhedonia, which is the lack of interest in pleasurable activities, is a core diagnostic criterion of depression which also contributes to reduced responsiveness to rewards (Huys et al., 2013), and alters the behavioral approach system (Henriques & Davidson, 2000; Pizzagalli et al., 2008). Findings in this study support this notion - neurobiologically, where we explored the impact of depression on the reward-related neural circuitry, which suggests that reduced sensitivity to rewards (i.e., reduced FN) among the low-SES is perhaps driven by their subclinical depressive symptoms.

FN in the current study is morphologically and topographical distribution is similar to what has been reported in studies with children who are experiencing depressive zhang(Bress et al., 2012; D. Zhang et al., 2020). FN is a variation of the FRN amplitude, which is largely driven by rewards and increase in negativity for losses (Warren & Holroyd, 2012; Yeung et al., 2005). Increasing evidence suggests that this increase in negativity for losses reflects the absence of positivity in loss trials, and FN is primarily characterized as an ongoing reward-related neural positivity (Baker & Holroyd, 2011; Foti et al., 2011; Holroyd et al., 2006; D. Zhang et al., 2020).

Consistent with previous studies (Bress et al., 2012), it was noticeable that participants depressive symptoms (i.e., CES-D scores) in the current study were associated with gains-related neural responses but not related to losses. Furthermore, socioeconomic status was also associated with depression and FN, which means a reduced FN amplitude was prominent among the low-SES. However, this noticeable SES-related neural sensitivity to rewards association was not prominent when depression was controlled for as a variable, which means that the link between SES and FN becomes non-existent. Taken together, results in the current study contribute to the existing FN-related literature suggesting that in the case of SES, reward sensitivity-related FN is driven by the presence of depressive symptoms.

When SES was considered, it is plausible to assume that the impact of depressive symptoms on the reward processing could also potentially be related to the heightened amygdala activity (to aversive stimuli) (E. Chen & Matthews, 2001; Herzberg & Gunnar, 2020; Javanbakht et al., 2015; Kuruvilla & Jacob, 2007; Muscatell et al., 2012). Changes in neurobiological responses among the low-SES are also in direct correlation with the early life stress model (Bick & Nelson, 2016). According to the early life stress model, prolonged exposure to early life stresses (associated with poverty, low income during the developmental stage) can activate physiological stress-related responses dysregulating homeostasis due to the accumulation of adversities (such as malnutrition, neighborhood violence, maltreatment, etc. (Evans, 2004) which can contribute to variabilities in reward processing (Sheridan & McLaughlin, 2014). However, the pattern of reward-related brain functions is associated with the type and the timing of the stress event. For instance, chronic exposure to poverty correlated with increased salience to reward (also heightened striatal activity), and socially deprived groups exhibiting an adaptive pattern of avoiding unpredictable losses (Evans, 2004; Herzberg & Gunnar, 2020). Our sample in the current study may have been clinically depressed, which has not been diagnosed in this study. Therefore, it is impossible to determine whether the participants suffered from depressive disorders or perhaps other co-morbid disorders. Further studies are required when conducting SES-related studies to differentiate whether FN is associated similarly with those who are clinically depressed.

Because most of the existing studies related to reward processing involve children, further studies are required to confirm our results whether a similar pattern exists across different age groups in low-SES. It is, however, important to emphasize that feedback used in the current does not reflect participants performance and cannot be used to interpret as such performance. Because studies that employ performance-based feedback has shown different results. For instance, the feedback-related error-related negativity (fERN) a performance-related ERP component which elicits at around 250 ms post-feedback, reflects when performance is worse than expected (Gehring & Willoughby, 2002; Luu et al., 2003). fERN negativity has been associated with vulnerable individuals who experience severe poverty (Herzberg & Gunnar, 2020), addictions and alcohol dependencies in which feedback-related neural responses are linked to individual performance and expectations. Findings in such studies are also consistent with existing non-performance-based FN studies (reduced FN among the depressed participants), although there are several studies that have found opposite results. Not SES-related, Tucker et al. (2003) had found a positive association between FN and depression when performance-based feedback processing was used in such studies. However, studies that employ purely reward and non-reward options are more specific towards identifying reward sensitivity which contributes to often different results than those with performance-based studies.

Given the association with depression, FN has the potential to become a biomarker of SES-related reward sensitivity when depression is considered. Yet, it is important to emphasize that the FN method cannot be directly associated with the effects specific to gains and losses responses, nor with clinical populations, which can be considered as limitations. If gains and losses are evaluated separately, there are potential artifactual issues (Luck, 2014). Therefore, the difference wave approach is suitable to overcome this artifactual problem by removing possible effects that can change gain and losses outcomes via experimental manipulations.

Due to the lack of SES-related FN studies, it is impossible to make a causal judgment that the SES *per se* can impact the sensitivity to reward feedback. However, the existing body of evidence related to depression and SES perhaps can shed light on the findings in this study. It is possible that reduced FN in the low-SES group was due to the presence of depressive symptoms compared to their affluent peers, because mood fluctuations has been found to be linked with blunted FN (Foti & Hajcak, 2010), which indicates a reward-related state-dependency response and are more prominent with those who have a history of depression (Foti et al., 2011). Future studies should evaluate whether the FN effect is a consequence of depression (clinical and subclinical) or persist independent of the SES status.

## AUTHOR CONTRIBUTIONS

HP: conceptualization, data collection, data analysis, manuscript writing, funding acquisition. RK and KS: supervision, review of the study design, analysis, revising the manuscript, funding acquisition.

## ACKNOWLEDGMENTS

The authors would like to thank Monash University Malaysia Neurobusiness Lab staff members for providing us with the Cognionics EEG equipment and all those who collaborated in the data collection.

## References

Aarts, K., Vanderhasselt, M.-A., Otte, G., Baeken, C., & Pourtois, G. (2013). Electrical brain imaging reveals the expression and timing of altered error monitoring functions in major depression. Journal of Abnormal Psychology, 122(4), 939–950. https://doi.org/10.1037/a0034616

Adamkovič, M., & Martončik, M. (2017). A Review of Consequences of Poverty on Economic Decision-Making: A Hypothesized Model of a Cognitive Mechanism. Frontiers in Psychology, 8. https://doi.org/10.3389/fpsyg.2017.01784

Amtmann, D., Kim, J., Chung, H., Bamer, A. M., Askew, R. L., Wu, S., Cook, K. F., & Johnson, K. L. (2014). Comparing CESD-10, PHQ-9, and PROMIS depression instruments in individuals with multiple sclerosis. Rehabilitation Psychology, 59(2), 220–229. https://doi.org/10.1037/a0035919

Anisman, H., & Matheson, K. (2005). Stress, depression, and anhedonia: Caveats concerning animal models. Neuroscience & Biobehavioral Reviews, 29(4–5), 525–546. https://doi.org/10.1016/j.neubiorev.2005.03.007

Baker, T. E., & Holroyd, C. B. (2011). Dissociated roles of the anterior cingulate cortex in reward and conflict processing as revealed by the feedback error-related negativity and N200. Biological Psychology, 87(1), 25–34. https://doi.org/10.1016/j.biopsycho.2011.01.010

Balconi, M., Finocchiaro, R., & Canavesio, Y. (2015). Reward Sensitivity (Behavioral Activation System), Cognitive, and Metacognitive Control in Gambling Behavior: Evidences From Behavioral, Feedback-Related Negativity, and P300 Effect. The Journal of Neuropsychiatry and Clinical Neurosciences, 27(3), 219–227. https://doi.org/10.1176/appi.neuropsych.14070165

Balleine, B. W., Delgado, M. R., & Hikosaka, O. (2007). The Role of the Dorsal Striatum in Reward and Decision-Making. Journal of Neuroscience, 27(31), 8161–8165. https://doi.org/10.1523/JNEUROSCI.1554-07.2007

Bechara, A., Damasio, A. R., Damasio, H., & Anderson, S. W. (1994). Insensitivity to future consequences following damage to human prefrontal cortex. Cognition, 50(1–3), 7–15. https://doi.org/10.1016/0010-0277(94)90018-3

Berghorst, L. H., Bogdan, R., Frank, M. J., & Pizzagalli, D. A. (2013). Acute stress selectively reduces reward sensitivity. Frontiers in Human Neuroscience, 7. https://doi.org/10.3389/fnhum.2013.00133

Bick, J., & Nelson, C. A. (2016). Early Adverse Experiences and the Developing Brain. Neuropsychopharmacology, 41(1), 177–196. https://doi.org/10.1038/npp.2015.252

Brand, M., Recknor, E. C., Grabenhorst, F., & Bechara, A. (2007). Decisions under ambiguity and decisions under risk: Correlations with executive functions and comparisons of two different gambling tasks with implicit and explicit rules. Journal of Clinical and Experimental Neuropsychology, 29(1), 86–99. https://doi.org/10.1080/13803390500507196

Braver, T. S., Krug, M. K., Chiew, K. S., Kool, W., Westbrook, J. A., Clement, N. J., Adcock, R. A., Barch, D. M., Botvinick, M. M., Carver, C. S., Cools, R., Custers, R., Dickinson, A., Dweck, C. S., Fishbach, A., Gollwitzer, P. M., Hess, T. M., Isaacowitz, D. M., Mather, M., … Somerville, L. H. (2014). Mechanisms of motivation–cognition interaction: challenges and opportunities. Cognitive, Affective, & Behavioral Neuroscience, 14(2), 443–472. https://doi.org/10.3758/s13415-014-0300-0

Bress, J. N., & Hajcak, G. (2013). Self-report and behavioral measures of reward sensitivity predict the feedback negativity. Psychophysiology, 50(7), 610–616. https://doi.org/10.1111/psyp.12053

Bress, J. N., Meyer, A., & Proudfit, G. H. (2015). The stability of the feedback negativity and its relationship with depression during childhood and adolescence. Development and Psychopathology, 27(4pt1), 1285–1294. https://doi.org/10.1017/S0954579414001400

Bress, J. N., Smith, E., Foti, D., Klein, D. N., & Hajcak, G. (2012). Neural response to reward and depressive symptoms in late childhood to early adolescence. Biological Psychology, 89(1), 156–162. https://doi.org/10.1016/j.biopsycho.2011.10.004

Carlson, J. M., Foti, D., Mujica-Parodi, L. R., Harmon-Jones, E., & Hajcak, G. (2011). Ventral striatal and medial prefrontal BOLD activation is correlated with reward-related electrocortical activity: A combined ERP and fMRI study. NeuroImage, 57(4), 1608–1616. https://doi.org/10.1016/j.neuroimage.2011.05.037

Carvalho, L. S., Meier, S., & Wang, S. W. (2016). Poverty and Economic Decision-Making: Evidence from Changes in Financial Resources at Payday. American Economic Review, 106(2), 260–284. https://doi.org/10.1257/aer.20140481

Carver, C. S., & White, T. L. (1994). Behavioral inhibition, behavioral activation, and affective responses to impending reward and punishment: The BIS/BAS Scales. Journal of Personality and Social Psychology, 67(2), 319–333. https://doi.org/10.1037/0022-3514.67.2.319

Chen, C.-C., & Yin, S.-J. (2008). Alcohol abuse and related factors in Asia. International Review of Psychiatry, 20(5), 425–433. https://doi.org/10.1080/09540260802344075

Chen, E., & Matthews, K. A. (2001). Cognitive appraisal biases: An approach to understanding the relation between socioeconomic status and cardiovascular reactivity in children. Annals of Behavioral Medicine, 23(2), 101–111. https://doi.org/10.1207/S15324796ABM2302_4

Chen, E., Miller, G. E., Brody, G. H., & Lei, M. (2015). Neighborhood Poverty, College Attendance, and Diverging Profiles of Substance Use and Allostatic Load in Rural African American Youth. Clinical Psychological Science, 3(5), 675–685. https://doi.org/10.1177/2167702614546639

Chi, Y. M., Wang, Y., Wang, Y.-T., Jung, T.-P., Kerth, T., & Cao, Y. (2013). A Practical Mobile Dry EEG System for Human Computer Interfaces (pp. 649–655). https://doi.org/10.1007/978-3-642-39454-6_69

Christie, G. J., & Tata, M. S. (2009). Right frontal cortex generates reward-related theta-band oscillatory activity. NeuroImage, 48(2), 415–422. https://doi.org/10.1016/j.neuroimage.2009.06.076

Cohen, J. D., & Blum, K. I. (2002). Reward and Decision. Neuron, 36(2), 193–198. https://doi.org/10.1016/S0896-6273(02)00973-X

Cools, R., Nakamura, K., & Daw, N. D. (2011). Serotonin and Dopamine: Unifying Affective, Activational, and Decision Functions. Neuropsychopharmacology, 36(1), 98–113. https://doi.org/10.1038/npp.2010.121

Crane, N. A., Funkhouser, C. J., Burkhouse, K. L., Klumpp, H., Phan, K. L., & Shankman, S. A. (2021). Cannabis users demonstrate enhanced neural reactivity to reward: An event-related potential and time-frequency EEG study. Addictive Behaviors, 113, 106669. https://doi.org/10.1016/j.addbeh.2020.106669

Crone, E. A., & van der Molen, M. W. (2004). Developmental Changes in Real Life Decision Making: Performance on a Gambling Task Previously Shown to Depend on the Ventromedial Prefrontal Cortex. Developmental Neuropsychology, 25(3), 251–279. https://doi.org/10.1207/s15326942dn2503_2

Cui, J., Chen, Y., Wang, Y., Shum, D. H. K., & Chan, R. C. K. (2013). Neural correlates of uncertain decision making: ERP evidence from the Iowa Gambling Task. Frontiers in Human Neuroscience, 7. https://doi.org/10.3389/fnhum.2013.00776

Dearing, E., Berry, D., & Zaslow, M. (n.d.). Poverty During Early Childhood. In Blackwell Handbook of Early Childhood Development (pp. 399–423). Blackwell Publishing Ltd. https://doi.org/10.1002/9780470757703.ch20

Delorme, A., & Makeig, S. (2004). EEGLAB: an open source toolbox for analysis of single-trial EEG dynamics including independent component analysis. Journal of Neuroscience Methods, 134(1), 9–21. https://doi.org/10.1016/j.jneumeth.2003.10.009

Dufford, A. J., Kim, P., & Evans, G. W. (2020). The impact of childhood poverty on brain health: Emerging evidence from neuroimaging across the lifespan (pp. 77–105). https://doi.org/10.1016/bs.irn.2019.12.001

Dunning, J. P., & Hajcak, G. (2007). Error-related negativities elicited by monetary loss and cues that predict loss. NeuroReport, 18(17), 1875–1878. https://doi.org/10.1097/WNR.0b013e3282f0d50b

Eamon, M. K., & Zuehl, R. M. (2001). Maternal depression and physical punishment as mediators of the effect of poverty on socioemotional problems of children in single-mother families. American Journal of Orthopsychiatry, 71(2), 218–226. https://doi.org/10.1037/0002-9432.71.2.218

Ellis, B. J., Figueredo, A. J., Brumbach, B. H., & Schlomer, G. L. (2009). Fundamental Dimensions of Environmental Risk. Human Nature, 20(2), 204–268. https://doi.org/10.1007/s12110-009-9063-7

Evans, G. W. (2004). The Environment of Childhood Poverty. American Psychologist, 59(2), 77–92. https://doi.org/10.1037/0003-066X.59.2.77

Flouri, E., Ruddy, A., & Midouhas, E. (2017). Maternal depression and trajectories of child internalizing and externalizing problems: the roles of child decision making and working memory. Psychological Medicine, 47(6), 1138–1148. https://doi.org/10.1017/S0033291716003226

Foti, D., & Hajcak, G. (2010). State sadness reduces neural sensitivity to nonrewards versus rewards. NeuroReport, 21(2), 143–147. https://doi.org/10.1097/WNR.0b013e3283356448

Foti, D., Kotov, R., Klein, D. N., & Hajcak, G. (2011). Abnormal Neural Sensitivity to Monetary Gains Versus Losses Among Adolescents at Risk for Depression. Journal of Abnormal Child Psychology, 39(7), 913–924. https://doi.org/10.1007/s10802-011-9503-9

Frank, M. J., & Claus, E. D. (2006). Anatomy of a decision: Striato-orbitofrontal interactions in reinforcement learning, decision making, and reversal. Psychological Review, 113(2), 300–326. https://doi.org/10.1037/0033-295X.113.2.300

Galván, M., Uauy, R., Corvalán, C., López-Rodríguez, G., & Kain, J. (2013). Determinants of Cognitive Development of Low SES Children in Chile: A Post-transitional Country with Rising Childhood Obesity Rates. Maternal and Child Health Journal, 17(7), 1243–1251. https://doi.org/10.1007/s10995-012-1121-9

Garg, A., Toy, S., Tripodis, Y., Cook, J., & Cordella, N. (2015). Influence of Maternal Depression on Household Food Insecurity for Low-Income Families. Academic Pediatrics, 15(3), 305–310. https://doi.org/10.1016/j.acap.2014.10.002

Garrido-Chaves, R., Perez-Alarcón, M., Perez, V., Hidalgo, V., Pulopulos, M. M., & Salvador, A. (2021). FRN and P3 during the Iowa gambling task: The importance of gender. Psychophysiology, 58(3). https://doi.org/10.1111/psyp.13734

Gehring, W. J., & Willoughby, A. R. (2002). The Medial Frontal Cortex and the Rapid Processing of Monetary Gains and Losses. Science, 295(5563), 2279–2282. https://doi.org/10.1126/science.1066893

Gianaros, P. J., & Hackman, D. (2013). Contributions of Neuroscience to the Study of Socioeconomic Health Disparities. Psychosomatic Medicine, 75(7), 610–615. https://doi.org/10.1097/PSY.0b013e3182a5f9c1

Gonzalez, M. Z., Allen, J. P., & Coan, J. A. (2016). Lower neighborhood quality in adolescence predicts higher mesolimbic sensitivity to reward anticipation in adulthood. Developmental Cognitive Neuroscience, 22, 48–57. https://doi.org/10.1016/j.dcn.2016.10.003

Haber, S. N., & Knutson, B. (2010). The Reward Circuit: Linking Primate Anatomy and Human Imaging. Neuropsychopharmacology, 35(1), 4–26. https://doi.org/10.1038/npp.2009.129

Hajcak, G., Moser, J. S., Holroyd, C. B., & Simons, R. F. (2006). The feedback-related negativity reflects the binary evaluation of good versus bad outcomes. Biological Psychology, 71(2), 148–154. https://doi.org/10.1016/j.biopsycho.2005.04.001

Heflin, C. M., & Iceland, J. (2009). Poverty, Material Hardship, and Depression. Social Science Quarterly, 90(5), 1051–1071. https://doi.org/10.1111/j.1540-6237.2009.00645.x

Henriques, J. B., & Davidson, R. J. (2000). Decreased responsiveness to reward in depression. Cognition & Emotion, 14(5), 711–724. https://doi.org/10.1080/02699930050117684

Herzberg, M. P., & Gunnar, M. R. (2020). Early life stress and brain function: Activity and connectivity associated with processing emotion and reward. NeuroImage, 209, 116493. https://doi.org/10.1016/j.neuroimage.2019.116493

Holroyd, C. B., Hajcak, G., & Larsen, J. T. (2006). The good, the bad and the neutral: Electrophysiological responses to feedback stimuli. Brain Research, 1105(1), 93–101. https://doi.org/10.1016/j.brainres.2005.12.015

Huys, Q. J., Pizzagalli, D. A., Bogdan, R., & Dayan, P. (2013). Mapping anhedonia onto reinforcement learning: a behavioural meta-analysis. Biology of Mood & Anxiety Disorders, 3(1), 12. https://doi.org/10.1186/2045-5380-3-12

Javanbakht, A., King, A. P., Evans, G. W., Swain, J. E., Angstadt, M., Phan, K. L., & Liberzon, I. (2015). Childhood Poverty Predicts Adult Amygdala and Frontal Activity and Connectivity in Response to Emotional Faces. Frontiers in Behavioral Neuroscience, 9. https://doi.org/10.3389/fnbeh.2015.00154

Jung, T. P., Makeig, S., Humphries, C., Lee, T. W., McKeown, M. J., Iragui, V., & Sejnowski, T. J. (2000). Removing electroencephalographic artifacts by blind source separation. Psychophysiology, 37(2), 163–178.

Keren, H., O’Callaghan, G., Vidal-Ribas, P., Buzzell, G. A., Brotman, M. A., Leibenluft, E., Pan, P. M., Meffert, L., Kaiser, A., Wolke, S., Pine, D. S., & Stringaris, A. (2018). Reward Processing in Depression: A Conceptual and Meta-Analytic Review Across fMRI and EEG Studies. American Journal of Psychiatry, 175(11), 1111–1120. https://doi.org/10.1176/appi.ajp.2018.17101124

Kim, P., Evans, G. W., Angstadt, M., Ho, S. S., Sripada, C. S., Swain, J. E., Liberzon, I., & Phan, K. L. (2013). Effects of childhood poverty and chronic stress on emotion regulatory brain function in adulthood. Proceedings of the National Academy of Sciences, 110(46), 18442–18447. https://doi.org/10.1073/pnas.1308240110

Kim, P., Neuendorf, C., Bianco, H., & Evans, G. W. (2016). Exposure to Childhood Poverty and Mental Health Symptomatology in Adolescence: A Role of Coping Strategies. Stress and Health, 32(5), 494–502. https://doi.org/10.1002/smi.2646

KreuSSel, L., Hewig, J., Kretschmer, N., Hecht, H., Coles, M. G. H., & Miltner, W. H. R. (2012). The influence of the magnitude, probability, and valence of potential wins and losses on the amplitude of the feedback negativity. Psychophysiology, 49(2), 207–219. https://doi.org/10.1111/j.1469-8986.2011.01291.x

Kring, A. M., & Bachorowski, J.-A. (1999). Emotions and Psychopathology. Cognition & Emotion, 13(5), 575–599. https://doi.org/10.1080/026999399379195

Kuruvilla, A., & Jacob, K. S. (2007). Poverty, social stress & mental health. The Indian Journal of Medical Research, 126(4), 273–278.

Lam, R. W., Kennedy, S. H., McIntyre, R. S., & Khullar, A. (2014). Cognitive Dysfunction in Major Depressive Disorder: Effects on Psychosocial Functioning and Implications for Treatment. The Canadian Journal of Psychiatry, 59(12), 649–654. https://doi.org/10.1177/070674371405901206

Lansford, J. E., Godwin, J., Bornstein, M. H., Chang, L., Deater-Deckard, K., di Giunta, L., Dodge, K. A., Malone, P. S., Oburu, P., Pastorelli, C., Skinner, A. T., Sorbring, E., Steinberg, L., Tapanya, S., Alampay, L. P., Uribe Tirado, L. M., Al-Hassan, S. M., & Bacchini, D. (2017). Reward sensitivity, impulse control, and social cognition as mediators of the link between childhood family adversity and externalizing behavior in eight countries. Development and Psychopathology, 29(5), 1675–1688. https://doi.org/10.1017/S0954579417001328

LeMoult, J., & Gotlib, I. H. (2019). Depression: A cognitive perspective. Clinical Psychology Review, 69, 51–66. https://doi.org/10.1016/j.cpr.2018.06.008

LeMoult, J., Humphreys, K. L., Tracy, A., Hoffmeister, J.-A., Ip, E., & Gotlib, I. H. (2020). Meta-analysis: Exposure to Early Life Stress and Risk for Depression in Childhood and Adolescence. Journal of the American Academy of Child & Adolescent Psychiatry, 59(7), 842–855. https://doi.org/10.1016/j.jaac.2019.10.011

Letourneau, N. L., Duffett-Leger, L., Levac, L., Watson, B., & Young-Morris, C. (2013). Socioeconomic Status and Child Development. Journal of Emotional and Behavioral Disorders, 21(3), 211–224. https://doi.org/10.1177/1063426611421007

Lopez-Calderon, J., & Luck, S. J. (2014). ERPLAB: an open-source toolbox for the analysis of event-related potentials. Frontiers in Human Neuroscience, 8. https://doi.org/10.3389/fnhum.2014.00213

Lorant, V. (2003). Socioeconomic Inequalities in Depression: A Meta-Analysis. American Journal of Epidemiology, 157(2), 98–112. https://doi.org/10.1093/aje/kwf182

Luck, S. (2014). An Introduction to the Event-Related Potential Technique, Second Edition | The MIT Press (Second). Bradford Books. https://mitpress.mit.edu/books/introduction-event-related-potential-technique-second-edition

Lund, C., de Silva, M., Plagerson, S., Cooper, S., Chisholm, D., Das, J., Knapp, M., & Patel, V. (2011). Poverty and mental disorders: breaking the cycle in low-income and middle-income countries. The Lancet, 378(9801), 1502–1514. https://doi.org/10.1016/S0140-6736(11)60754-X

Luu, P., Tucker, D. M., Derryberry, D., Reed, M., & Poulsen, C. (2003). Electrophysiological Responses to Errors and Feedback in the Process of Action Regulation. Psychological Science, 14(1), 47–53. https://doi.org/10.1111/1467-9280.01417

MacCallum, R. C., Zhang, S., Preacher, K. J., & Rucker, D. D. (2002). On the practice of dichotomization of quantitative variables. Psychological Methods, 7(1), 19–40. https://doi.org/10.1037/1082-989X.7.1.19

Mazlan, N. H., & Ahmad, A. (2014). Validation of the Malay-translated version of the Center for Epidemiological Study—Depression Scale (CES-D). ASEAN Journal of Psychiatry, 15(1), 54–65. https://psycnet.apa.org/record/2014-39533-007

McLaughlin, K. A., Sheridan, M. A., & Lambert, H. K. (2014). Childhood adversity and neural development: Deprivation and threat as distinct dimensions of early experience. Neuroscience & Biobehavioral Reviews, 47, 578–591. https://doi.org/10.1016/j.neubiorev.2014.10.012

McLoyd, V. C. (1998). Socioeconomic disadvantage and child development. American Psychologist, 53(2), 185–204. https://doi.org/10.1037/0003-066X.53.2.185

Miltner, W. H. R., Braun, C. H., & Coles, M. G. H. (1997). Event-Related Brain Potentials Following Incorrect Feedback in a Time-Estimation Task: Evidence for a “Generic” Neural System for Error Detection. Journal of Cognitive Neuroscience, 9(6), 788–798. https://doi.org/10.1162/jocn.1997.9.6.788

Moser, J. S., & Simons, R. F. (2009). The neural consequences of flip-flopping: The feedback-related negativity and salience of reward prediction. Psychophysiology, 46(2), 313–320. https://doi.org/10.1111/j.1469-8986.2008.00760.x

Mueller, E. M., Pechtel, P., Cohen, A. L., Douglas, S. R., & Pizzagalli, D. A. (2015). POTENTIATED PROCESSING OF NEGATIVE FEEDBACK IN DEPRESSION IS ATTENUATED BY ANHEDONIA. Depression and Anxiety, 32(4), 296–305. https://doi.org/10.1002/da.22338

Mullen, T. R., Kothe, C. A. E., Chi, Y. M., Ojeda, A., Kerth, T., Makeig, S., Jung, T.-P., & Cauwenberghs, G. (2015). Real-time neuroimaging and cognitive monitoring using wearable dry EEG. IEEE Transactions on Biomedical Engineering, 62(11), 2553–2567. https://doi.org/10.1109/TBME.2015.2481482

Muscatell, K. A., Morelli, S. A., Falk, E. B., Way, B. M., Pfeifer, J. H., Galinsky, A. D., Lieberman, M. D., Dapretto, M., & Eisenberger, N. I. (2012). Social status modulates neural activity in the mentalizing network. NeuroImage, 60(3), 1771–1777. https://doi.org/10.1016/j.neuroimage.2012.01.080

Mussel, P., Reiter, A. M. F., Osinsky, R., & Hewig, J. (2015). State- and trait-greed, its impact on risky decision-making and underlying neural mechanisms. Social Neuroscience, 10(2), 126–134. https://doi.org/10.1080/17470919.2014.965340

Noble, K. G., Farah, M. J., & McCandliss, B. D. (2006). Socioeconomic background modulates cognition–achievement relationships in reading. Cognitive Development, 21(3), 349–368. https://doi.org/10.1016/j.cogdev.2006.01.007

Noble, K. G., Houston, S. M., Brito, N. H., Bartsch, H., Kan, E., Kuperman, J. M., Akshoomoff, N., Amaral, D. G., Bloss, C. S., Libiger, O., Schork, N. J., Murray, S. S., Casey, B. J., Chang, L., Ernst, T. M., Frazier, J. A., Gruen, J. R., Kennedy, D. N., van Zijl, P., … Sowell, E. R. (2015). Family income, parental education and brain structure in children and adolescents. Nature Neuroscience, 18(5), 773–778. https://doi.org/10.1038/nn.3983

Nusslock, R., & Miller, G. E. (2016). Early-Life Adversity and Physical and Emotional Health Across the Lifespan: A Neuroimmune Network Hypothesis. Biological Psychiatry, 80(1), 23–32. https://doi.org/10.1016/j.biopsych.2015.05.017

Olvet, D. M., Klein, D. N., & Hajcak, G. (2010). Depression symptom severity and error-related brain activity. Psychiatry Research, 179(1), 30–37. https://doi.org/10.1016/j.psychres.2010.06.008

Ong, Q., Theseira, W., & Ng, I. Y. H. (2019). Reducing debt improves psychological functioning and changes decision-making in the poor. Proceedings of the National Academy of Sciences, 116(15), 7244–7249. https://doi.org/10.1073/pnas.1810901116

Osinsky, R., Ulrich, N., Mussel, P., Feser, L., Gunawardena, A., & Hewig, J. (2017). The Feedback-related Negativity Reflects the Combination of Instantaneous and Long-term Values of Decision Outcomes. Journal of Cognitive Neuroscience, 29(3), 424–434. https://doi.org/10.1162/jocn_a_01055

Patel, V. (2001). Depression in developing countries: lessons from Zimbabwe. BMJ, 322(7284), 482–484. https://doi.org/10.1136/bmj.322.7284.482

Payne, B. K., Brown-Iannuzzi, J. L., & Hannay, J. W. (2017). Economic inequality increases risk taking. Proceedings of the National Academy of Sciences, 114(18), 4643–4648. https://doi.org/10.1073/pnas.1616453114

Peirce, J., Gray, J. R., Simpson, S., MacAskill, M., Höchenberger, R., Sogo, H., Kastman, E., & Lindeløv, J. K. (2019). PsychoPy2: Experiments in behavior made easy. Behavior Research Methods, 51(1), 195–203. https://doi.org/10.3758/s13428-018-01193-y

Perera-W.A., H. (2022). A Brief Guide for Using Cognionics HD-72 Wireless EEG System: Noninvasive Method of Measuring the Brain’s Electric Fields in Psychophysiological Research. PsyArXiv Preprints. https://doi.org/10.31234/OSF.IO/VCYFB

Perera-W.A., H., Salehuddin, K., Khairudin, R., & Schaefer, A. (2021). The Relationship Between Socioeconomic Status and Scalp Event-Related Potentials: A Systematic Review. Frontiers in Human Neuroscience, 15. https://doi.org/10.3389/fnhum.2021.601489

Pizzagalli, D. A., Iosifescu, D., Hallett, L. A., Ratner, K. G., & Fava, M. (2008). Reduced hedonic capacity in major depressive disorder: Evidence from a probabilistic reward task. Journal of Psychiatric Research, 43(1), 76–87. https://doi.org/10.1016/j.jpsychires.2008.03.001

Potts, G. F., Martin, L. E., Burton, P., & Montague, P. R. (2006). When Things Are Better or Worse than Expected: The Medial Frontal Cortex and the Allocation of Processing Resources. Journal of Cognitive Neuroscience, 18(7), 1112–1119. https://doi.org/10.1162/jocn.2006.18.7.1112

Radloff, L. S. (1977). The CES-D Scale. Applied Psychological Measurement, 1(3), 385–401. https://doi.org/10.1177/014662167700100306

Ridley, M., Rao, G., Schilbach, F., & Patel, V. (2020). Poverty, depression, and anxiety: Causal evidence and mechanisms. Science, 370(6522). https://doi.org/10.1126/science.aay0214

Romens, S. E., Casement, M. D., McAloon, R., Keenan, K., Hipwell, A. E., Guyer, A. E., & Forbes, E. E. (2015). Adolescent girls’ neural response to reward mediates the relation between childhood financial disadvantage and depression. Journal of Child Psychology and Psychiatry, 56(11), 1177–1184. https://doi.org/10.1111/jcpp.12410

Ruchensky, J. R., Bauer, E. A., & MacNamara, A. (2020). Intolerance of uncertainty, depression and the error-related negativity. International Journal of Psychophysiology, 153, 45–52. https://doi.org/10.1016/j.ijpsycho.2020.04.015

Sambrook, T. D., & Goslin, J. (2014). Mediofrontal event-related potentials in response to positive, negative and unsigned prediction errors. Neuropsychologia, 61, 1–10. https://doi.org/10.1016/j.neuropsychologia.2014.06.004

Santesso, D. L., Steele, K. T., Bogdan, R., Holmes, A. J., Deveney, Christen M., Meites, T. M., & Pizzagalli, D. A. (2008). Enhanced negative feedback responses in remitted depression. NeuroReport, 19(10), 1045–1048. https://doi.org/10.1097/WNR.0b013e3283036e73

Schultz, W. (2006). Behavioral Theories and the Neurophysiology of Reward. Annual Review of Psychology, 57(1), 87–115. https://doi.org/10.1146/annurev.psych.56.091103.070229

Shafer, A. B. (2006). Meta-analysis of the factor structures of four depression questionnaires: Beck, CES-D, Hamilton, and Zung. Journal of Clinical Psychology, 62(1), 123–146. https://doi.org/10.1002/jclp.20213

Shankman, S. A., Klein, D. N., Tenke, C. E., & Bruder, G. E. (2007). Reward sensitivity in depression: A biobehavioral study. Journal of Abnormal Psychology, 116(1), 95–104. https://doi.org/10.1037/0021-843X.116.1.95

Shankman, S. A., Nelson, B. D., Sarapas, C., Robison-Andrew, E. J., Campbell, M. L., Altman, S. E., McGowan, S. K., Katz, A. C., & Gorka, S. M. (2013). A psychophysiological investigation of threat and reward sensitivity in individuals with panic disorder and/or major depressive disorder. Journal of Abnormal Psychology, 122(2), 322–338. https://doi.org/10.1037/a0030747

Sheridan, M. A., & McLaughlin, K. A. (2014). Dimensions of early experience and neural development: deprivation and threat. Trends in Cognitive Sciences, 18(11), 580–585. https://doi.org/10.1016/j.tics.2014.09.001

Tang, Y., Zhang, X., Simmonite, M., Li, H., Zhang, T., Guo, Q., Li, C., Fang, Y., Xu, Y., & Wang, J. (2013). Hyperactivity within an extensive cortical distribution associated with excessive sensitivity in error processing in unmedicated depression: A combined event-related potential and sLORETA study. International Journal of Psychophysiology, 90(2), 282–289. https://doi.org/10.1016/j.ijpsycho.2013.09.001

Tomarken, A. J., Dichter, G. S., Garber, J., & Simien, C. (2004). Resting frontal brain activity: linkages to maternal depression and socioeconomic status among adolescents. Biological Psychology, 67(1–2), 77–102. https://doi.org/10.1016/j.biopsycho.2004.03.011

Toplak, M. E., Sorge, G. B., Benoit, A., West, R. F., & Stanovich, K. E. (2010). Decision-making and cognitive abilities: A review of associations between Iowa Gambling Task performance, executive functions, and intelligence. Clinical Psychology Review, 30(5), 562–581. https://doi.org/10.1016/j.cpr.2010.04.002

Tucker, D. M., Luu, P., Frishkoff, G., Quiring, J., & Poulsen, C. (2003). Frontolimbic Response to Negative Feedback in Clinical Depression. Journal of Abnormal Psychology, 112(4), 667–678. https://doi.org/10.1037/0021-843X.112.4.667

Tversky, A., & Kahneman, D. (1982). Evidential impact of base rates. In Judgment under Uncertainty (pp. 153–160). Cambridge University Press. https://doi.org/10.1017/CBO9780511809477.011

Tversky, A., & Kahneman, D. (1992). Advances in prospect theory: Cumulative representation of uncertainty. Journal of Risk and Uncertainty, 5(4), 297–323. https://doi.org/10.1007/BF00122574

van der Maas, M. (2016). Problem gambling, anxiety and poverty: an examination of the relationship between poor mental health and gambling problems across socioeconomic status. International Gambling Studies, 16(2), 281–295. https://doi.org/10.1080/14459795.2016.1172651

Warren, C. M., & Holroyd, C. B. (2012). The Impact of Deliberative Strategy Dissociates ERP Components Related to Conflict Processing vs. Reinforcement Learning. Frontiers in Neuroscience, 6. https://doi.org/10.3389/fnins.2012.00043

Webb, C. A., Auerbach, R. P., Bondy, E., Stanton, C. H., Foti, D., & Pizzagalli, D. A. (2017). Abnormal neural responses to feedback in depressed adolescents. Journal of Abnormal Psychology, 126(1), 19–31. https://doi.org/10.1037/abn0000228

Yaple, Z. A., & Yu, R. (2020). Functional and Structural Brain Correlates of Socioeconomic Status. Cerebral Cortex, 30(1), 181–196. https://doi.org/10.1093/cercor/bhz080

Yeung, N., Holroyd, C. B., & Cohen, J. D. (2005). ERP Correlates of Feedback and Reward Processing in the Presence and Absence of Response Choice. Cerebral Cortex, 15(5), 535–544. https://doi.org/10.1093/cercor/bhh153

Zhang, D., Shen, J., Bi, R., Zhang, Y., Zhou, F., Feng, C., & Gu, R. (2020). Differentiating the abnormalities of social and monetary reward processing associated with depressive symptoms. Psychological Medicine, 1–15. https://doi.org/10.1017/S0033291720003967

Zhang, W.-N., Chang, S.-H., Guo, L.-Y., Zhang, K.-L., & Wang, J. (2013). The neural correlates of reward-related processing in major depressive disorder: A meta-analysis of functional magnetic resonance imaging studies. Journal of Affective Disorders, 151(2), 531–539. https://doi.org/10.1016/j.jad.2013.06.039

